# Ethanol stress induces siRNA-mediated epimutations and phenotypic variability in fission yeast

**DOI:** 10.64898/2026.10.04.756277

**Authors:** Mariana Villalba de la Peña, Peter Sarkies

## Abstract

Epigenetic information is required for the maintenance of gene expression states through cell division. Some epigenetic information can also persist through generations, raising the possibility that epigenetic differences may arise in populations and contribute to evolutionary processes such as drift and natural selection. Evaluating this requires characterising the properties of such epimutations, so that their rate, stability and genome-wide distribution can be assessed. Recent experiments in many organisms have demonstrated that epimutations occur, but have limited stability relative to DNA sequence mutations. However, the extent to which epimutations contribute to evolution is still unclear, in part because it is not clear whether epimutations respond dynamically to environmental conditions. Here, we use the fission yeast *S. pombe*, in which evolutionarily relevant epimutations have been observed, but not yet systematically characterised, to test whether a physiologically relevant stress, ethanol exposure, leads to changes in the properties of epimutations. Using laboratory evolution experiments with repeated population bottlenecks to minimise selection, we show that epimutations associated with changes in small interfering (si)RNAs occur at a greater rate in ethanol. Ethanol-induced epimutations are enriched at transposable element loci and correlate with changes in transposon expression. Ethanol exposure also increases gene expression noise genome-wide and variability in growth rate; importantly, both these increases are suppressed in mutants lacking siRNAs. Together, these results demonstrate that stress can alter the generation of heritable epigenetic information, and suggest that siRNAs may have a role in promoting phenotypic variability in response to environmental stress.

## Introduction

The traditional view of evolution considers the transmission of genetic material to be the primary hereditary system driving evolutionary change. However, this perspective has long been debated due to substantial evidence suggesting that evolutionary processes can also occur through alternative, non-genetic hereditary systems, such as epigenetic inheritance (Laland et al., 2015; Stajic and Jansen, 2021).

A fundamental principle of the modern evolutionary synthesis is the random generation of genetic mutations. This principle has long been supported by foundational genetic research (Luria and Delbrück, 1943), which indicates that, although mutation rates may vary depending on environmental conditions or genetic context (Villalba de la Peña et al., 2023; Wei et al., 2022), beneficial mutations arise randomly and are not inherently predisposed to occur in response to environmental stressors (Svensson and Berger, 2019). Epimutations, however, may accumulate differently and could potentially be triggered in stress-response genes following exposure to stressful conditions (Ashe et al., 2021).

Studying the properties of epimutations requires examining epimutation accumulation under conditions in which the effects of selection are minimized. In an approach derived from classical attempts to study mutation rates in DNA (genetic mutations) in whole organisms, ‘epimutation accumulation’ experiments have been performed in plants (Van Der Graaf et al., 2015; Schmitz et al., 2011; Becker et al., 2011), the nematode worm *C. elegans* (Beltran et al., 2020; Wilson et al., 2023; Fallet et al., 2023) and the filamentous fungus *N. crassa* (de la Peña et al., 2026). These experiments have converged on the general picture that epimutations occur at much greater rates than genetic mutations, but have reduced stability (Ganem and Sarkies, 2026). However, these experiments were largely performed under benign conditions, leaving the issue of whether epimutations change in either rate, stability or genome-wide distribution under stress unclear. Changes in the distribution or stability of epimutations under stress could be very important in understanding their potential to contribute to evolutionary adaptation: whilst epimutations that occur in stable, benign conditions are short-lived and thus unlikely to reach high frequencies within populations, epimutations that emerge specifically in response to stress could promote fixation of stable, beneficial genetic mutations (Webster and Phillips, 2024).

Here, we set out to investigate how the properties of epimutations change in response to stressful environments. We reasoned that using a physiological stressor would maximize the possibility that the organism will have evolved responses to the stressor, potentially including adaptive changes in epimutation rate. We therefore decided to use the fission yeast *S. pombe* as our study system for these experiments.

*S. pombe* is a well-characterized model organism that shares several epigenetic features with higher eukaryotes, and the mechanisms underlying heterochromatin formation in this species are particularly well defined. In *S. pombe*, heterochromatin formation is mediated in part by RNA interference (RNAi) through the RNA-induced transcriptional silencing (RITS) complex. In this pathway, the RNA-dependent RNA polymerase Rdp1 converts transcripts, primarily those originating from repetitive regions, into double-stranded RNA, which is subsequently processed into small interfering RNAs (siRNAs) by Dicer (Dcr1). These siRNAs provide sequence specificity by associating with Ago1 and directing the RITS complex towards complementary genomic regions. The RNAi machinery interacts with the H3K9 methyltransferase Clr4, which subsequently promotes the recruitment of heterochromatin-associated proteins (Swi6 and Chp proteins) (Cam et al., 2005). RNAi-dependent heterochromatin is particularly associated with genomic regions such as centromeres, subtelomeres, ribosomal DNA repeats, and the mating-type locus (Cam et al., 2005). Crucially, previous studies have shown that epimutations occur in *S. pombe*, driven by changes in heterochromatin that lead to silencing of specific loci and result in resistance to drugs (Torres-Garcia et al., 2020). However, these were performed under conditions of strong selection for drug resistance. Systematic characterization of *S. pombe* epimutations in the absence of selection has not yet been performed. Moreover, the ability of *S. pombe* to tolerate moderate exposure to ethanol stress enabled us to apply a physiologically relevant stressor continuously throughout an epimutation accumulation experiment. We therefore performed epimutation accumulation over several bottlenecks to reduce selection, and studied the properties of epimutation in the presence or absence of ethanol. We showed that epimutations arise frequently, have limited duration and are strongly enriched for transposable element-derived loci. However, ethanol significantly increased the rate at which epimutations occur genome-wide. Indeed, this was associated with increased transcriptional heterogeneity and increased heterogeneity in growth rates between different clonal lineages. Remarkably, deletion of the siRNA machinery suppressed both the increase in gene expression variability and the increase in growth rate heterogeneity due to ethanol exposure, supporting a link between epimutations and phenotypic heterogeneity. Taken together, our results demonstrate that ethanol exposure increases epimutation rates, thus demonstrating stress-induced generation of epigenetic variation, leading to increased phenotypic heterogeneity which gives some lineages the potential for increased survival.

## Results

### Ethanol stress induces distinct transcriptomic and siRNA responses

We performed an epimutation accumulation experiment (eMA) in which four lines derived from a single ancestor were propagated either in YES medium (two control lines) or in YES medium with 4% ethanol (two stress lines). The lines were propagated for 10 transfers, bottle-necking the population at each transfer to minimize the action of natural selection (Figure 1A). We sequenced siRNAs at all ten transfers and mRNA from the ancestor and at transfers five and ten. Raw siRNA and mRNA reads were aligned to the reference *S. pombe* genome using Bowtie and HISAT2, respectively (see Methods). Aligned siRNA reads were filtered by size to retain reads between 22-26 bp and subsequently grouped into 1-kb windows.

**Figure 1:**
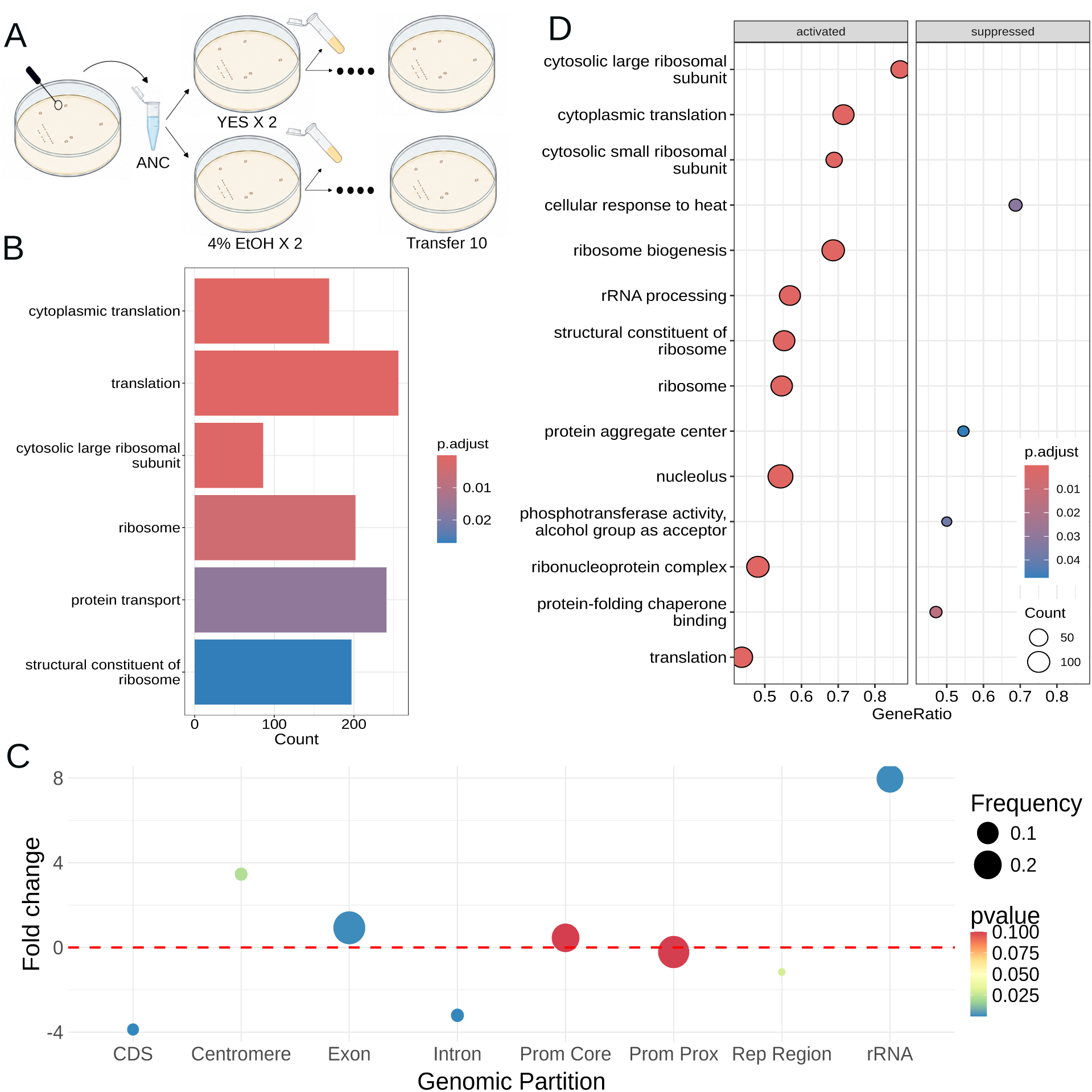
A) Experimental design. Cells from a single colony were diluted and propagated in four lines and two different treatments. In every transfer only a few cells from a single colony were used to propagate on the next plate and cultured in liquid media to obtain enough material for RNA extractions. **B)** Results from the over-representation analysis (ORA) showing the pathways where siRNAs were enriched in the ancestor. **C)** Genomic regions showing enrichment for siRNAs in the ancestor. **D)** Shows pathways overrepresented for differential gene expression (mRNA) upon ethanol exposure. **Alt text:** Four-panel figure showing the experimental design and molecular responses across the eMA lines. Panel A depicts the propagation of four lines under two treatments through repeated single-colony transfers, followed by RNA extraction. Panel B is a bar plot showing that siRNAs in the ancestor are enriched for genes involved in protein synthesis. Panel C is a bubble plot showing that siRNA-enriched regions in the ancestor are enriched at rRNA loci and the centromere. Panel D is a bubble plot showing that protein-synthesis-associated genes are enriched for gene-expression changes under ethanol conditions.

We first characterized the overall siRNA profile. siRNAs in the ancestral population were enriched in genes associated with translation, ribosomal activity, and protein transport (Figure 1B). Across all lines, siRNAs occurred more frequently than expected by chance in centromeric regions, exons, and ribosomal RNA genes, whereas they were depleted in coding sequences (CDS), introns, promoters, and repeat regions (Figure 1C).

To characterize the global molecular response of *S. pombe* to ethanol stress, we identified differentially expressed genes (DEGs) and differentially enriched siRNA regions (DERs) between the control and ethanol treatments. We identified 6,074 DEGs between treatments, indicating an extensive transcriptional response to ethanol stress. Genes up-regulated under ethanol stress were predominantly associated with ribosomal and translation-related processes (Figure 1D). In contrast, only four biological processes were significantly enriched among genes down-regulated under ethanol stress, including processes associated with the heat-stress response and protein folding.

Changes in siRNA enrichment were considerably less extensive than changes in gene expression. We identified 115 DERs between treatments, most of which overlapped lncRNA loci, centromeric repeats, or rRNA loci. However, over-representation analysis did not identify any significantly enriched biological pathways among DER-associated genes. The transcriptional and siRNA responses showed little overlap. Only 27 genes were associated with both DEGs and DERs, the majority of which were lncRNA and rRNA genes, with only a small proportion corresponding to protein-coding genes. Importantly, this overlap was not greater than expected by chance (Fisher’s exact test, *P >* 0.05). Thus, although ethanol stress induces extensive changes in gene expression and a more limited redistribution of siRNAs, we found no evidence that ethanol-responsive changes in siRNA enrichment are preferentially associated with transcriptionally responsive genes.

### Ethanol stress increases the epimutation rate

To characterize epimutations, we calculated a modified Z-score from the fold change in normalized siRNA counts between the ancestor and each eMA line at each transfer. Windows with a modified Z-score above 3 or below *−*3 were classified as having increased or decreased siRNA abundance relative to the ancestor, respectively, as this maximised agreement between technical replicates. Windows with modified Z-scores between *−*3 and 3 were classified as unchanged (see Methods and Figure S1B). Thus, each window at each transfer was assigned one of three states: 1 for increased siRNA abundance, *−*1 for decreased abundance, and 0 for no change relative to the ancestor.

We defined a transition as any change in state between two consecutive transfers. An epimuta-tion was defined as a change in state that persisted for at least two consecutive transfers following a transition (Figure S1C). We use transfers, rather than generations, as the temporal unit because a generation in *S. pombe* corresponds to a single cell division. However, we reasoned that if an epigenetic change remains stable for at least two consecutive transfers, it is likely that this change was maintained throughout the intervening cell cycles.

We first examined how epigenetic divergence among samples from the same lineage changed over successive transfers. If changes in siRNA states accumulate and persist over time, samples separated by a greater number of transfers are expected to become increasingly divergent. A linear regression model was used to assess whether correlation changed as a function of the number of generations. Under 4% ethanol, siRNA divergence increased significantly with transfer distance (*F*_(1,88)_ = 7.72, *p <* 0.01). In the control treatment, this relationship was weaker and did not reach statistical significance (*F*_(1,79)_ = 3.24, *p* = 0.07, Figure 2A). In both treatments, the observed divergence was not predominantly driven by centromeres and telomeres (ethanol: *F*_(1,88)_ = 0.52, *p >* 0.05, control: *F*_(1,79)_ = 2.35, *p >* 0.05; Figure S2).

**Figure 2:**
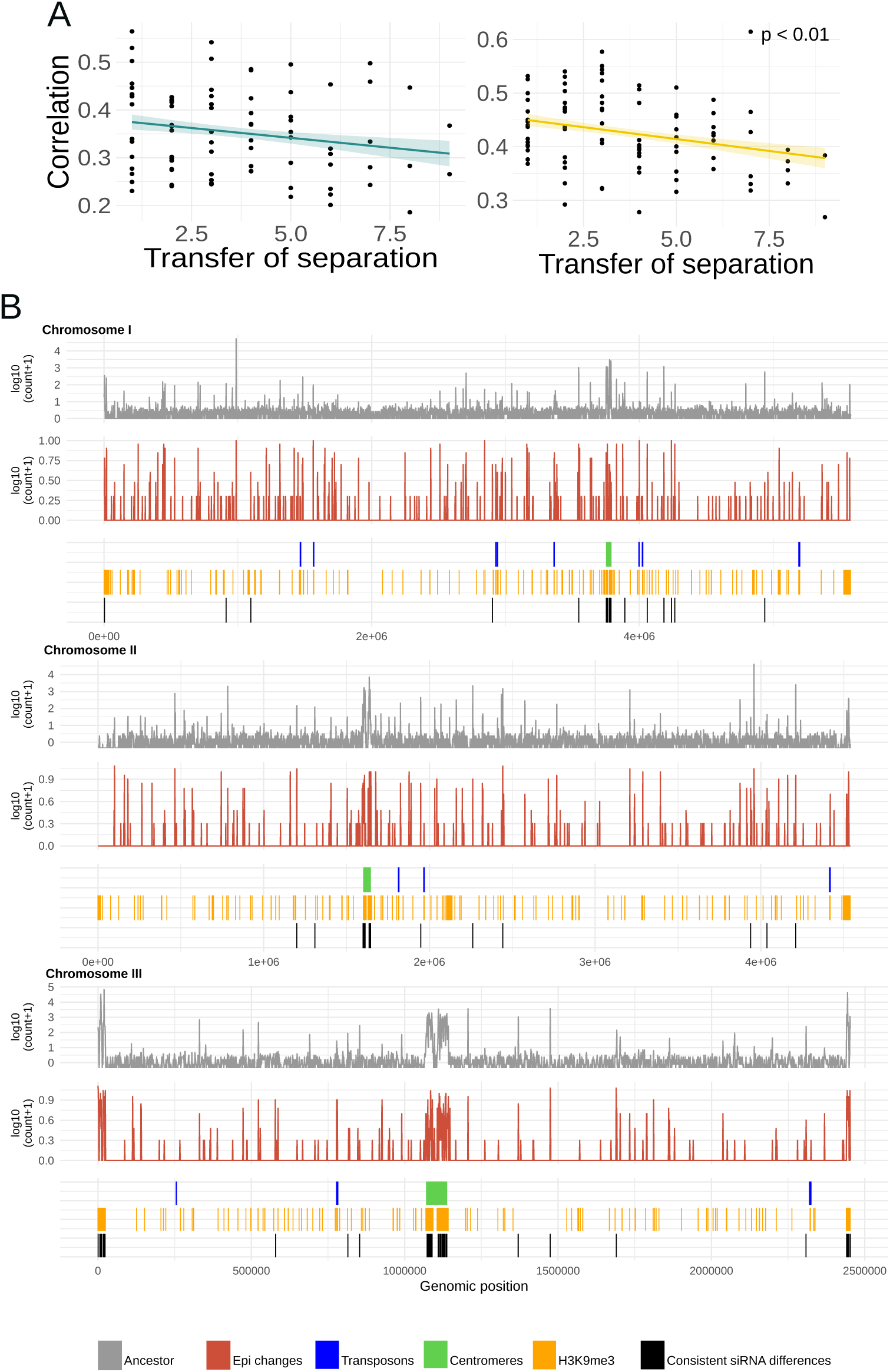
A) Divergence of epigenetic changes in the control (green) and ethanol-treated (yellow) lines. **B)** Genome-wide distribution of genomic and epigenetic features. The distribution of siRNAs along the genome, in the ancestor, is shown in grey, epimutations in red, transposon positions in blue, centromeric regions in green, and regions marked by H3K9me in orange. Publicly available *S. pombe* H3K9me ChIP-seq data were downloaded from the NCBI Sequence Read Archive (SRA), using run accessions SRR4449335–SRR4449338 and SRR4449455–SRR4449458. Black regions represent genomic regions where siRNAs were consistently changed under ethanol stress. **Alt text:** Two panel figure showing characterisation of epimutations under control conditions and ethanol stress. **A** shows that correlation between epimutations decreases with time within lineages suggesting longitudinal inheritance. **B** shows the genome-wide distribution of epimutations demonstrating enrichment within heterochromatic regions but clear epimutation hotspots outside of heterochromatin.

To assess the stability of epimutations we performed a survival analysis (Wilson et al., 2023). We found that most epigenetic changes were short-lived (Figure S3): only 1.3 – 1.6% of the epimutations persist through all 10 transfers, though as above we note that 10 transfers corresponds to several cell divisions (Figure S3 A).

Finally, we tested whether ethanol stress affected the frequency at which transitions and epimutations occurred. In total, we identified 2,332 transitions under ethanol stress compared with 1,925 in the control, and 1,059 epimutations under ethanol stress compared with 884 in the control. Both transitions (*SE* = 0.06, *z* = 3.30, *p <* 0.001, TableS1) and epimutations (*SE* = 0.08, *z* = 2.31, *p <* 0.05, Table S1) were significantly more frequent under ethanol stress than under control conditions (Figure S3 B). Together, these results indicate that ethanol stress increased the rate at which changes in siRNA states arise.

### Epimutations target transposable elements under ethanol stress

Epimutations were not solely located in heterochromatin regions. Although epimutations accumulated in centromeres (*p* = 9.999*e^−^*^05^) and repetitive regions (*p* = 9.999*e^−^*^05^) more than expected by chance, they also accumulated at a high frequency in exons and promoter regions.

Epimutations did not accumulate at the genes where siRNAs were originally enriched, or in the DERs between ethanol and control. We performed an over-representation analysis to examine if the genes that overlapped with epimutated windows were involved in certain molecular pathways. In both treatments there was an enrichment in functions related to transposon activity and the genes that were listed within the enriched pathways were transposons (supplementary file). However, the higher numbers of epimutations and transitions in ethanol seem to be concentrated in fewer genes (19 genes) compared with control (43 genes). While in ethanol most of the genes in epimutated windows were restricted mainly to TEs, in control conditions the genes were less constrained, spreading over other genes and pathways.

To investigate whether epimutations are associated with changes in gene expression, we compared siRNA and mRNA fold changes within epimutated genomic windows. Epimutated windows showed heterogeneous patterns, with increases or decreases in mRNA expression occurring alongside either increased or decreased siRNA abundance. However, transposon overlapping regions were significantly enriched among windows exhibiting high mRNA expression and low siRNA abundance compared with the expectation under random assignment of these patterns (*p <* 0.001; 10,000 permutations; Figure 3B; Table S2). Together, these results show that epimutations are not randomly distributed across the genome but preferentially occur in transposable elements. Moreover, the enrichment of transposons among epimutated regions with reduced siRNA abundance and increased mRNA expression suggests that epimutations at these loci are associated with loss of siRNA-mediated repression and increased transposon transcription.

**Figure 3:**
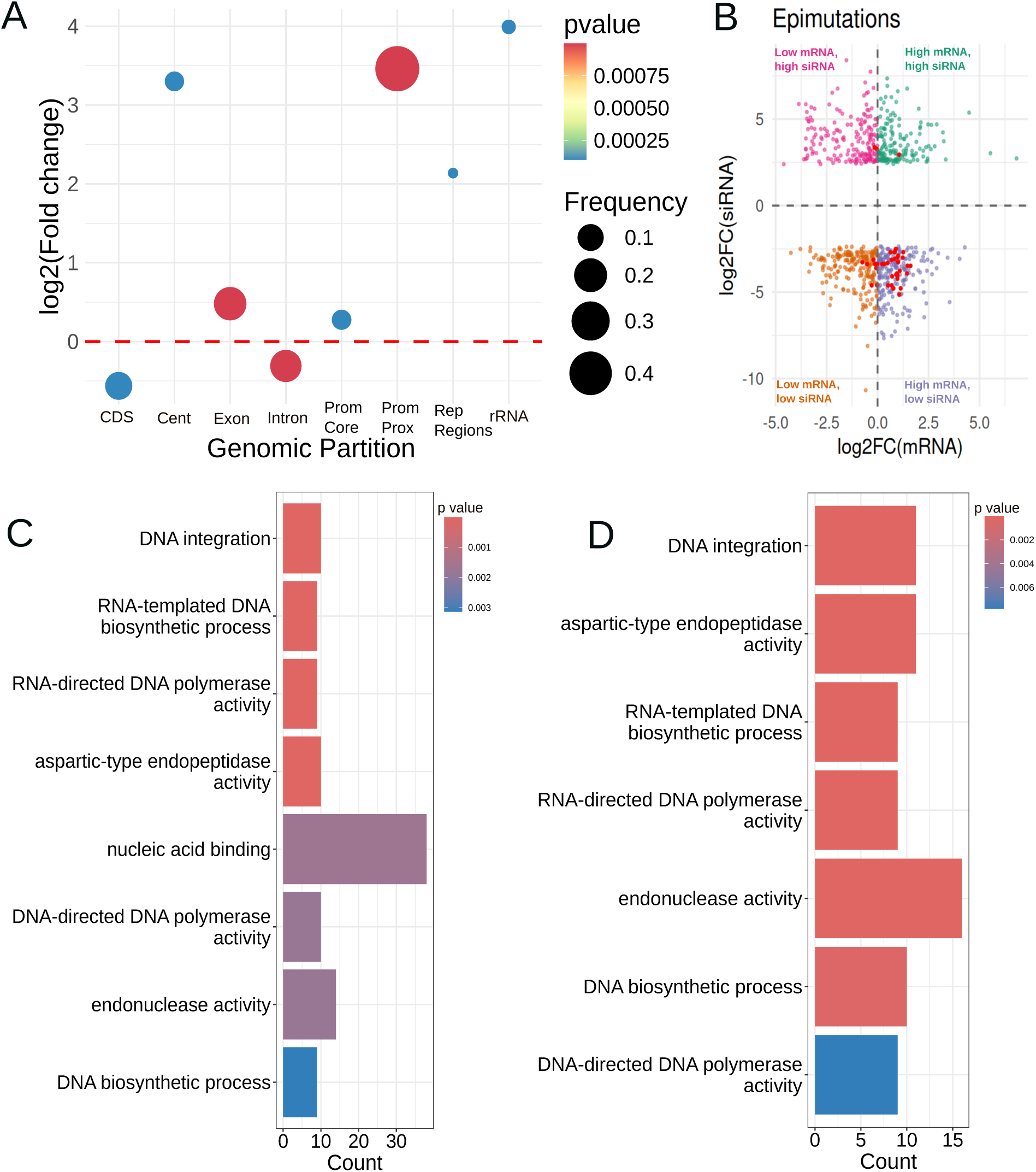
A) Distribution of epimutations across different genomic features. **B)** Correlation between siRNA counts and gene mRNA counts within each genomic window. Colors represent different relationships between gene mRNA and siRNA abundance, depending on whether mRNA and siRNA counts are high or low. Red dots indicate windows containing transposons. **C)** and **D)** Results of the overrepresentation analysis of epimutations across genomic features in the control (**C**) and ethanol-treated (**D**) lines. **Alt text:** Characterisation of siRNA-mediated epimutations in *S. pombe*. Four panel figure. **A** is a bubble plot showing that epimutations are enriched in the centromere, the promoter and repeat regions, as well as rRNA loci (compare to Figure 1C). **B** is a scatter plot showing the fold change in siRNAs vs change in mRNAs for epimutated regions. Epimutations at TEs are highlighted and show enrichment for negative correlation between mRNA and siRNA changes, indicating repression of TEs by siRNAs. **C** and **D** show enrichment for functions related to TEs in genes affected by epimutations, in both control conditions (**C**) and under ethanol stress (**D**).

### siRNAs promote phenotypic and transcriptional variability under ethanol stress

Given the increased epimutation rate and the preferential accumulation of epimutations in transposable elements under ethanol stress, we hypothesized that siRNA-mediated epigenetic variability could contribute to phenotypic diversification and potentially facilitate adaptation to stressful conditions. To investigate this possibility, we measured growth rates over five transfers in YES medium supplemented with 6% ethanol and in control YES medium. We compared three strains: wild type (WT), Δ*ago*1, and Δ*rdp*1, the latter two lacking key components of the siRNA pathway. We found no evidence of adaptation over the five transfers in the WT strains, as growth rates did not consistently increase over time (Figure 4 A). Both mutants showed a trend towards increased growth rates over time, which was similar in control and ethanol conditions (Figure 4 A). However, phenotypic variability differed significantly among strains and treatments. Under ethanol stress, the WT showed the greatest variability in growth rate, which was 5.9-fold higher than that observed for the WT under control conditions. Although variability also increased in the Δ*ago*1 and Δ*rdp*1 strains under ethanol stress, it was only 1.8– and 2.5-fold higher, respectively, than that of the WT under control conditions. Thus, disruption of the siRNA pathway reduced the phenotypic variability associated with stress (Figure 4 B).

**Figure 4:**
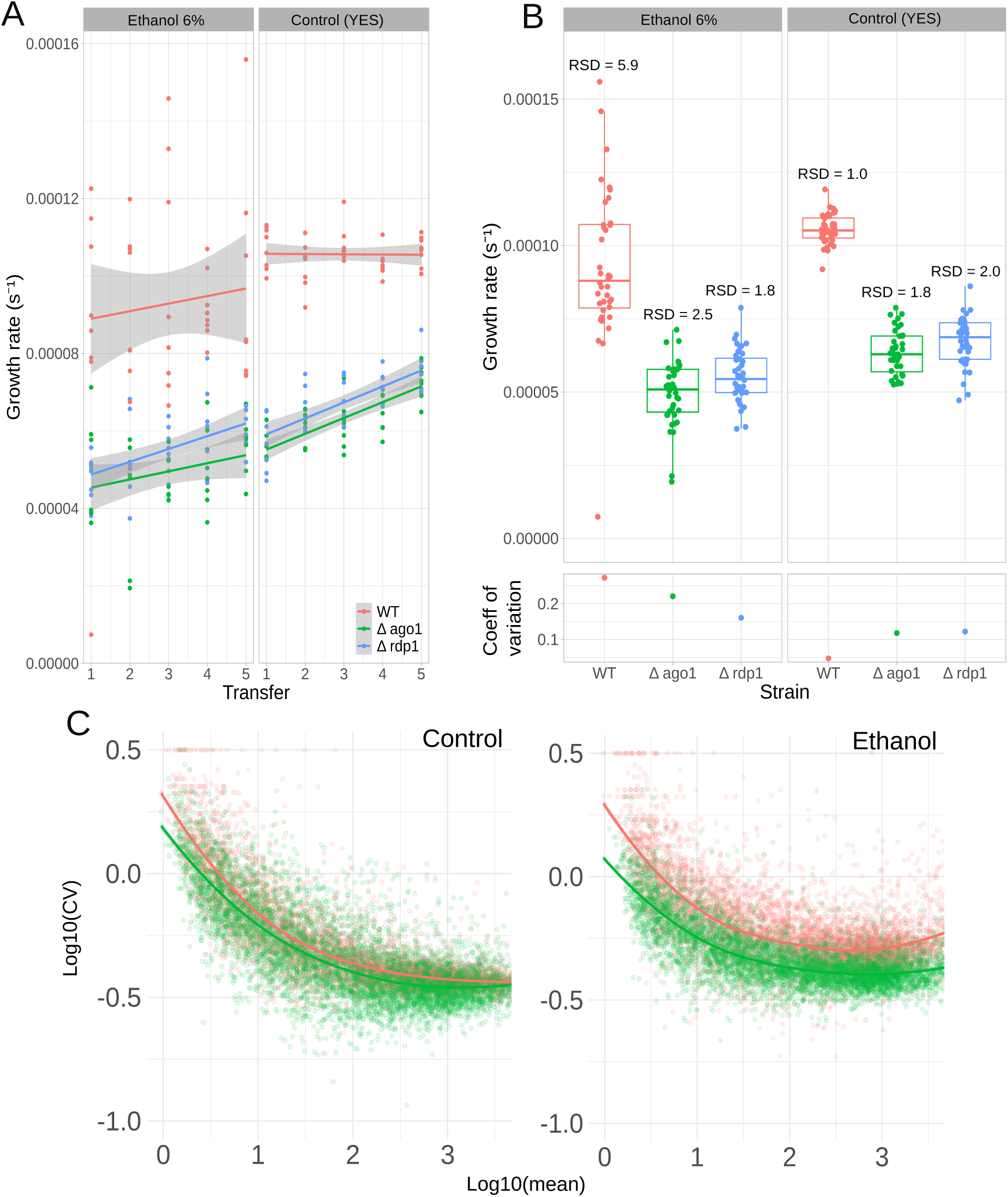
A) Growth rates of the WT and the Δ*ago1* and Δ*rdp*1 mutants across five transfers in liquid YES medium (control) and YES medium supplemented with 6% ethanol. **B)** The relative standard deviation (RSD; coefficient of variation) is shown to illustrate the variability in growth rate for each strain across transfers. **C)** LOESS regression between the mean expression of each gene across the 10 replicates and its coefficient of variation. The WT is shown in pink and the Δ*ago1* mutant in green. **Alt text:** Analysis of growth rate changes over experimental evolution in either ethanol or control conditions. **A** is a scatter plot showing how growth rate changes over multiple transfers. WT show no significant increase in growth rate in either ethanol or control conditions; mutants lacking siRNAs (Δ*ago1* and Δ*rdp1*) show comparable increases in growth rate in either control or ethanol conditions. **B** is a box plot showing that there is a large increase in growth rate variability in WT under ethanol exposure, but variability is largely similar for both mutants between ethanol and control conditions. **C** shows the relationship between gene expression noise (measured as coefficient of variation) and mean expression, with a LOESS fit. The curves for WT and Δ*ago1* are similar in control conditions, but WT shows higher gene expression noise in ethanol.

To determine whether this increased phenotypic variability was accompanied by greater variability in gene expression, we performed an additional mutation accumulation experiment. Ten independent lines of the WT and Δ*ago*1 strains were propagated for five transfers under either 4% ethanol or control conditions, after which mRNA was sequenced. To distinguish technical from biological variability, total RNA samples were spiked with 0.5% *C. elegans* total RNA. After accounting for technical noise (see Methods), we calculated the coefficient of variation (CV) in expression for each gene across replicate lines.

Consistent with the phenotypic results, the Δ*ago*1 strain exhibited significantly lower gene expression variability than the WT under both control and ethanol conditions. Importantly, the difference between the strains was substantially greater under ethanol stress: the WT showed approximately 9.6% higher expression CV than Δ*ago*1 under control conditions, compared with approximately 29.8% higher CV under ethanol stress. Both strain and treatment therefore had significant effects on gene-expression variability(Figure 4C; Table S1).

A similar pattern was observed when the analysis was restricted to transposable elements. As expected from the loss of siRNA-mediated silencing, mean transposon expression was higher in the Δ*ago*1 mutant. In contrast, variability in transposon expression was consistently greater in the WT. Thus, loss of Ago1 increases the average expression of transposable elements while simultaneously reducing variation in their expression among independent lines.

Together, these results indicate that an intact siRNA pathway is associated with increased phenotypic and transcriptional variability, particularly under ethanol stress. Although we did not detect increased adaptation over the timescale of our experiment, the enhanced variability observed in the WT provides a potential mechanism for generating phenotypic diversity under stressful conditions. Combined with the increased accumulation of siRNA epimutations under ethanol stress, these findings are consistent with a model in which siRNA-mediated epigenetic variation contributes to a potential bet-hedging strategy by increasing the range of phenotypes available to a population under environmental stress.

## Discussion

Understanding the role of epigenetic inheritance in adaptation and evolution requires a clear understanding of the properties of epimutation and a comparison to the properties of genetic mutations. Here we examined a fundamental property of epimutation: does it respond to environmental conditions? Using ethanol exposure in *S. pombe* as a model we show significantly increased epimutation rate in response to an environmentally realistic stress. Importantly, we provide evidence for a link between increased epimutation rate and growth rate heterogeneity, which is reduced by removing components of the RNA interference machinery. Below, we interpret these findings in the context of previous investigation of epimutations across systems, and speculate on the implications for evolution driven by epigenetic change.

### The properties of epimutations in *S. pombe* under benign conditions

Although the existence of heritable epimutations has been established at single gene loci in *S. pombe* (Torres-Garcia et al., 2020), our study provides the first systematic study of their properties in the absence of selection. One important property of epimutations is their stability. Epimutation accumulation experiments in *C. elegans* (Beltran et al., 2020), *Arabidopsis* (e.g. (Van Der Graaf et al., 2015; Yao et al., 2021), rice (Zheng et al., 2017) and the filamentous fungus *N. crassa* (de la Peña et al., 2026) have shown that epimutations are more frequent than genetic mutations but short-lived. It is interesting that despite a wide variety in the molecular mechanisms underpinning epimutations in these systems, the median time over which an epimutation persists is usually less than 10 generations. Here we showed that epimutations in *S. pombe* were short-lived, with a median duration of 2 transfers under benign conditions. It is worth noting that in our experiment ‘transfer’ is not directly equivalent to ‘generation’ in a cultured nematode or plant species: in between transfers there will be a small number of cell divisions as the colony grows on a new plate. This means that one transfer probably corresponds to as many as 10 mitotic divisions. Nevertheless, it seems clear that *S. pombe*, despite again having a distinct system of establishment and maintenance of epimutations, has comparable epimutation duration to other organisms. It is possible that the limited stability of epimutations may itself be a property that has evolved in order to maximize their benefit, perhaps by preventing permanent ‘accidental’ silencing of important genomic regions. However, it is also possible that, despite the differences in molecular mechanisms, epigenetic inheritance systems are fundamentally less able to specify precise states (Yao et al., 2021; Radman-Livaja et al., 2011), thus can never achieve the same degree of fidelity enjoyed by DNA replication.

Another feature of *S. pombe* epimutations that aligns with previous studies is their elevated rate compared to DNA sequence changes. Across the genome, we found an average of approximately 100 epimutations per transfer, whilst <1 DNA sequence mutation is the rate estimated from classical mutation accumulation experiments. Under a simple Markov model of epimutations, the high rate of epimutations is intrinsically linked to their limited duration (Ganem and Sarkies, 2026; Yao et al., 2021). Due to the differences in the way epimutations are defined, it is difficult to compare the precise number of epimutations between species; nevertheless the number is comparable to *C. elegans*, taking into account the smaller genome of *S. pombe*. It is important to note as above that the lack of ability to precisely determine ‘generations’ in the *S. pombe* epimutation lines affects the rate estimate, in particular because the short period of growth between bottlenecks imposes selection, which may cause detrimental epimutations to be preferentially lost thus underestimating the final epimutation rate. Overcoming this limitation would require assessment of epimutations genome-wide at a single cell level within a lineage, which is currently impossible.

Our experiments also studied the genome-wide distribution of epimutations. Again concordant with other species, epimutations showed a highly biased genome-wide distribution. Nevertheless, they did not simply congregate around regions generally established to be rich in siRNAs, namely centromeres, and did not exclusively overlap with heterochromatin, indicating increased plasticity relative to stable heterochromatic regions. Intriguingly the predominant location of epimutations was in transposable elements. In *S. pombe*, TEs constitute approximately 1.1% of the genome and are derived from long terminal repeat retrotransposons (Esnault and Levin, 2015). They comprise two closely related subfamilies, Tf1 and Tf2, which belong to the *gypsy* family. The reference strain contains 13 full-length copies of Tf2 elements but no full-length Tf1 elements, although Tf1 elements can be found in wild isolates (Esnault and Levin, 2015). Interestingly, previous studies have shown that Tf2 silencing is largely independent of siRNAs in the wild type, but that abundant siRNAs do form against Tf2 in the context of deletion of the RNA degradation machinery (Yamanaka et al., 2013). Our identification of transposable elements as a hotspot for epimutations suggests that siRNAs may be more transiently associated with TEs, hence appearing and disappearing in separate lines but not being abundant in bulk cultures. It is interesting that the majority of the epimutations we observed at TEs were losses of siRNAs, and corresponded to increased RNA levels. One possible explanation for this directional bias is that siRNAs assist in silencing TEs, and that this is under purifying selection. Thus when selection is relaxed, such as in epimutation accumulation experiments, reduced siRNAs and increased expression result. In this regard, it is interesting that we also showed that the entire loss of the siRNA pathway resulted in overall increased TE expression. However, it is also possible that this bias was simply a chance effect due to the ancestor of all the populations having a somewhat elevated level of siRNAs at these loci. Further experiments with larger numbers of lines would be needed to resolve this question.

### Stress induced epimutations and their implication for evolution

Our study is one of the few published attempts to explore how the properties of epimutation are affected by stress. We found that ethanol, a physiologically relevant stressor for fission yeast, led to an increased rate of epimutation but a decreased genomic range of epimutations, such that epimutations became even more heavily concentrated in TEs. There are two broad possible explanations for increased epimutation rate under ethanol exposure. One would be simply that stress weakens the stability of epigenetic inheritance, so that epimutations appear and disappear more rapidly, in the same way that many molecular systems fail under stress. Alternatively, increased epimutation might be an evolved response either to ethanol or to stress in general, perhaps as a bet-hedging strategy. In support of the possibility that increased epimutations might be an evolved response to ethanol, we showed that phenotypic variability was increased in response to ethanol exposure in WT, including the generation of some lines with improved survival. Indeed, we observed increased variability in expression of TEs in response to ethanol in WT. Variability in phenotype and TE expression was much less pronounced in mutants deficient for RNAi. Given the predominance of epimutations at TEs, this supports a direct link between epimutations and TE expression variability. In other organisms, TE expression changes have been linked to stressful conditions, and proposed to provide a survival advantage (Tossolini et al., 2025). How TE expression variability might influence variability in growth rate is an open question, though it is interesting to note that gene expression variability was also increased in WT but not mutant strains, and TE expression has pervasive influences on genome-wide gene expression, often through complex effects on gene networks (Fueyo et al., 2022). What might the relevance of these results be for the question of whether epimutations might influence the course of evolution? *S. pombe* provides one of the few model systems where an unambiguous example of laboratory evolution prompted by epimutations has been observed (Torres-Garcia et al., 2020). Our results establish that epimutations are not rare events, but occur frequently both under benign conditions and ethanol stress. We did not observe epimutations at the *hda-1* locus that drives drug resistance; however it is possible that the spectrum of induced epimutations might be dependent on the particular stress applied. It is also possible that the mechanism of *hda-1* epimutation, which is dependent on heterochromatin, is independent of siRNAs (Sorida and Murakami, 2020). Further research is needed to determine how different stressors, including toxic drug exposure, alter the epimutation accumulation pattern in the absence of selection. In addition, developing a more complete picture of epimutation accumulation patterns will require investigating other epigenetic mechanisms associated with heterochromatin formation, such as H3K9me. Understanding whether the epimutation landscape itself has evolved to best suit different stresses would provide strong evidence that epigenetic inheritance could play a key role in environmental adaptation and, ultimately, evolution.

## Methods

### Epimutation accumulation experiment

The ancestor of the eMA lines was a culture derived from a single colony of the WT strain MCW1975 (h-ade6-M210). To conserve the ancestor line a small fraction of these cells were cultured in 5 ml of YES liquid media and frozen. The remainder were spread on plates with YES agar to create two lines grown in YES media (control environment) and two lines grown in YES media with 4% ethanol (stress environment). Every three days, a single colony was randomly chosen, and transferred to the next plate. We repeated this process for ten transfers. During each transfer few cells were propagated in 5 ml liquid culture for two days to get enough cell material for a glycerol and a Trizol frozen stock, later used for RNA extraction.

### RNA extraction

We performed RNA extraction from the frozen trizol stocks as specified in (Roth et al., 2018). For the samples used to measure gene expression noise, we added 0.5% of *C. elegans* total RNA to all of the samples as an external spike to exclude technical noise. Total RNA was sent to Novogene for siRNA and mRNA library construction and sequencing. mRNA library preparation was via poly-A enrichment and paired-end sequencing was performed using NovaSeq sequencing platform. siRNA libraries were sequenced using the Illumina Sequencing SE50 platform.

### Characterization of epigenetic changes and epimutations

We selected siRNA reads that were between 21 and 27 nucleotides. We aligned them, using bowtie (version 1.3.1), to the *S. pombe* genome, updated on September 2025 in pombase. (https://www.pombase.org/monthly_releases/2025/pombase-2025-09-01/). RNA alignments were normalized using R package **DeSeq2** (version 1.48.2: (Love et al., 2014)). To identify epimutations, we followed the procedure described in (Wilson et al., 2023) with a few modifications to better describe the properties of siRNAs in *S. pombe*. We first divided the genome into non-overlapping 1kb windows. In each window we calculated the fold change in the DESeq-normalized siRNA level using the ancestor as a reference. Then we calculated a non-parametric modified Z-score (1) for each bin.

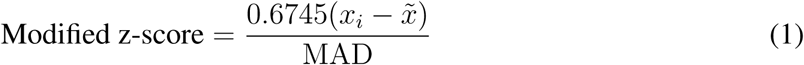

To determine a threshold to identify epimutations we plotted the change between the normalized siRNA count of the ancestor versus the normalized eMA-line, and fitted a LOESS curve to the scatter plot in the 10 samples that had technical replicates. Then we calculated the Z-scores by obtaining the difference of the observed fold change minus the average of the LOESS fit, divided by the standard deviation of the LOESS fit. We repeated this analysis with five different thresholds (1 to 5) and, we selected the threshold that showed more shared epimutations between the technical replicates. We used 10 samples with their technical replicates to establish the threshold (Figure S1A). L2T9 acted as an outlier so we removed this sample from the analysis. This analysis converged on a threshold of +-3; thus if the modified Z-score was above 3 we would assign a 1 to the window, if the score was below *−*3 we would assign a *−*1 to the window and if the score was between 3 and *−*3 we would assign a 0 to the window. We then retained this threshold for the remainder of the analyses to ensure consistency across datasets.

### Divergence and median survival time of epimutations

We used the three levels of epimutation data, encoded as *−*1, 0, and 1, as an input to calculate divergence, stability and number of epimutations and transitions. To address if epigenetic changes are accumulating randomly we calculated the divergence of the epigenetic changes using pariwise Spearman correlations between transfers from the same eMA-line. To calculate the median survival time of the epimutations we counted the number of consecutive time points in which the same epimutation state was maintained at each genomic region. Epimutations that were still present at the last sampled time point were considered censored. Persistence times were calculated separately for each lineage and then grouped into control and ethanol-treated conditions. Kaplan–Meier survival analysis was used to compare how long epimutations persisted between the two conditions. We performed this analysis using the R package **survival** (version 3.8-6; (Therneau, 2026)) and **survminer** (version 0.5.2; (Kassambara et al., 2026)).

To calculate the number of epimutations and transitions, we examined the epimutation state of each genomic window across consecutive transfers within each lineage. Only genomic windows showing at least one epigenetic change in any of the samples were included. Stable epimutations were defined as the same state (*−*1, 0, or 1) occurring for at least two consecutive transfers, and the number of stable events of each type was recorded. We also counted the number of times each genomic window changed epigenetic state between consecutive transfers.

### Growth rate

We grew 8 replicates of the WT, Δago1 and Δrdp1 in 5 ml of either YES media or YES media with 6% ethanol, for five transfers. After two days these were collected, 1 ml of the culture were transferred to the next culture tube and 1M cells were transferred to a 200ul well in a 96 well plate. We then measured OD600 every 10 minutes 200 times (for a total of 33 hours). This data was analyzed using the R package **growthCurver** (version 0.3.1; (Sprouffske and Wagner, 2016)) to obtain the growth rate.

### Gene expression analysis

mRNA reads were aligned to the reference genome using **HISAT2** (version 2.2.2), and gene-level read counts were obtained using **featureCounts** (version 2.1.1). Data normalization and differential gene expression analysis were performed using **DESeq2** (version 1.48.2; (Love et al., 2014)).

To measure gene expression noise, *C. elegans*-mapped reads were used as external spike-ins. Counts were normalized using the R package *RUVSeq* (version 1.42.0; (Risso et al., 2014)), first by upper-quartile normalization and then using RUVg with the spike-in genes and one factor of unwanted variation (*k* = 1). RLE plots and PCA were used to assess normalization (Figure S4). The normalized counts for *S. pombe* genes were then extracted for further analysis.

### Statistical Analysis

To model the number of epimutations and transitions we used a generalised linear mixed-effect model using the number of transitions or epimutations as the response variable:

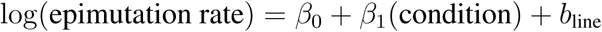

where *β*_0_ represents the intercept, *β*_1_ the effect of condition, and *b*_line_ the random effect associated with each experimental line.

We considered treatment, plus the number of transfers in that line as the fixed effect, and the line as the random factor, using the Poisson distribution. We checked for over dispersion in all of the models. We included the number of transfers as a fixed effect, and removed transfers that were outliers.

To determine whether the mutant strains exhibited greater gene expression noise compared with the WT, we calculated the coefficient of variation (CV) for each gene. We fitted three linear mixed-effects models using the log-transformed CV as the response variable. First, control and ethanol-treated samples were analyzed separately (2), with strain included as a fixed effect and gene ID as a random intercept to account for gene-specific variation. We then fitted a combined model including strain, treatment, and their interaction as fixed effects, with gene ID again included as a random factor (3). The strain-by-treatment interaction was used to assess whether the effect of strain on gene expression noise differed between control and ethanol conditions.

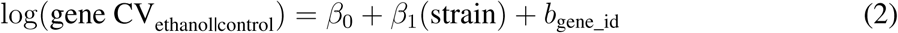

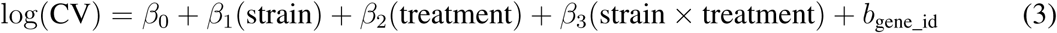

### Permutation tests

To examine the genomic distribution of siRNAs and epimutations, we used the R package **GenomicDistributions** (version 1.16.1; (Kupkova et al., 2022)) to determine their overlap with CDSs, centromeres, exons, introns, promoter core and proximal regions, repeat regions, and rRNAs. We then used **regioneR** (version 1.40.1; (Gel et al., 2016)) to perform 10,000 permutation tests, randomly redistributing the query regions across the genome and comparing the observed number of overlaps with the randomized distribution to determine whether each genomic feature was significantly enriched or depleted.

To assess the relationship between siRNA and mRNA levels, we calculated the fold change for each genomic window by comparing each eMA line at a given time point with the ancestor. Windows were classified as upregulated when the fold-change was greater than 0 and downregulated when it was below 0, and were subsequently assigned to four categories: up in both mRNA and siRNA, down in both, up in mRNA and down in siRNA, or down in mRNA and up in siRNA. As transposons predominantly overlapped windows with increased mRNA and decreased siRNA, we performed a permutation test in which quadrant assignments were randomized while controlling for sample identity, allowing us to determine whether the observed transposon overlap was greater than expected by chance.

### Data accessibility

The raw sequencing data generated in this study is deposited in NCBI under the accession number: PRJNA1535449

Scripts and phenotypic data is available in the Github repository https://github.com/villab04/siRNA-epimutations-ms

## Supplementary Figures

**Figure S1:**
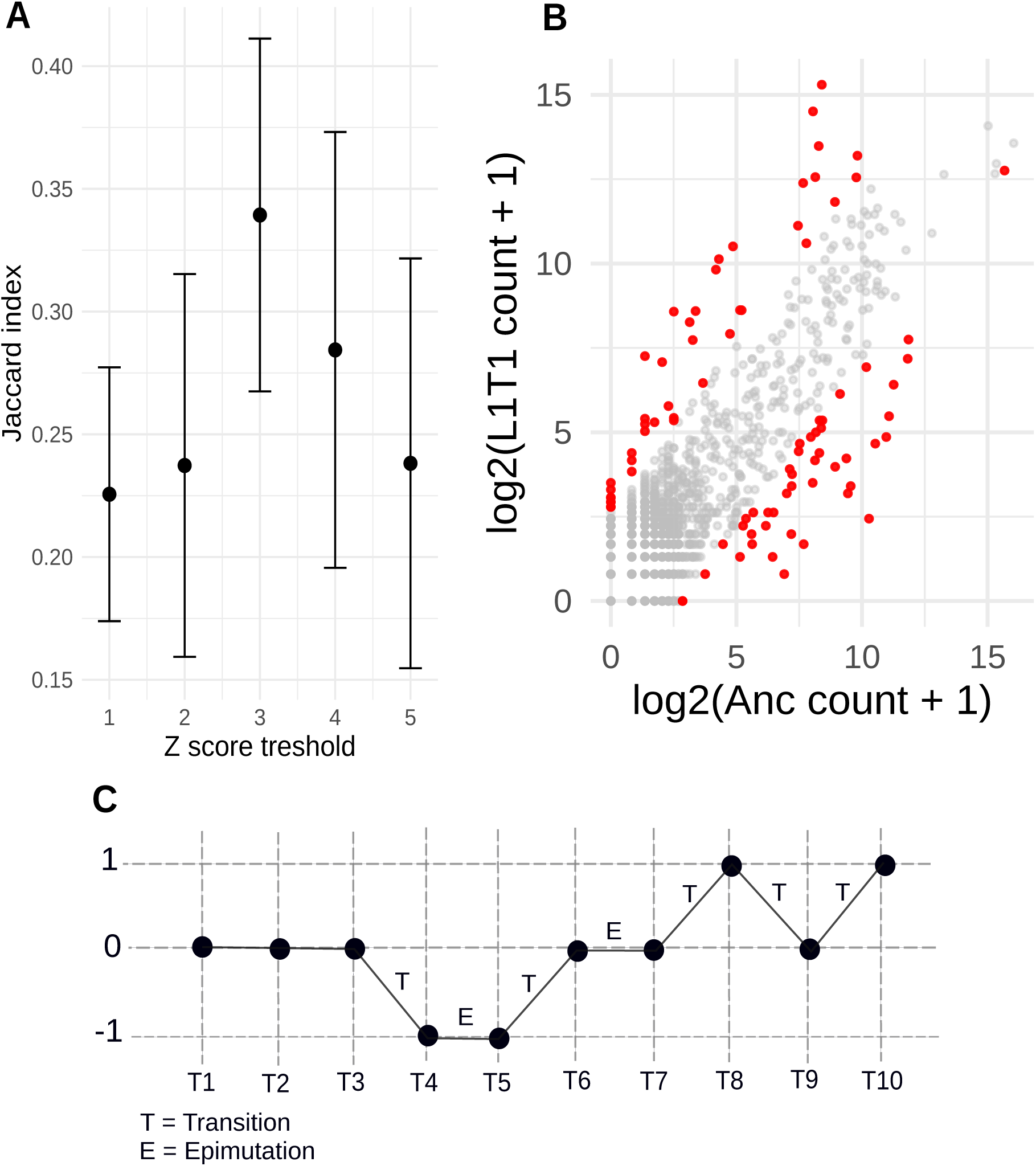
A) Jaccard index across Z-score thresholds. A Z-score threshold of 3 showed the highest Jaccard index, indicating that epimutations called in technical replicates were most similar at this threshold. **B)** Example using eMA line L1 at transfer 1. The upper red dots represent windows with a modified Z-score above 3, which were assigned a value of 1, whereas the lower red dots represent windows with a modified Z-score below *−*3, which were assigned a value of *−*1. Thus, the red dots represent windows that differ from the ancestral state. **C)** Example of how epimutations were called across the ten transfers of a single eMA line. Each change in state among 1, 0, and *−*1 was considered a transition (T). Following a transition, a state that remained stable for at least two transfers was considered an epimutation (E).

**Figure S2:**
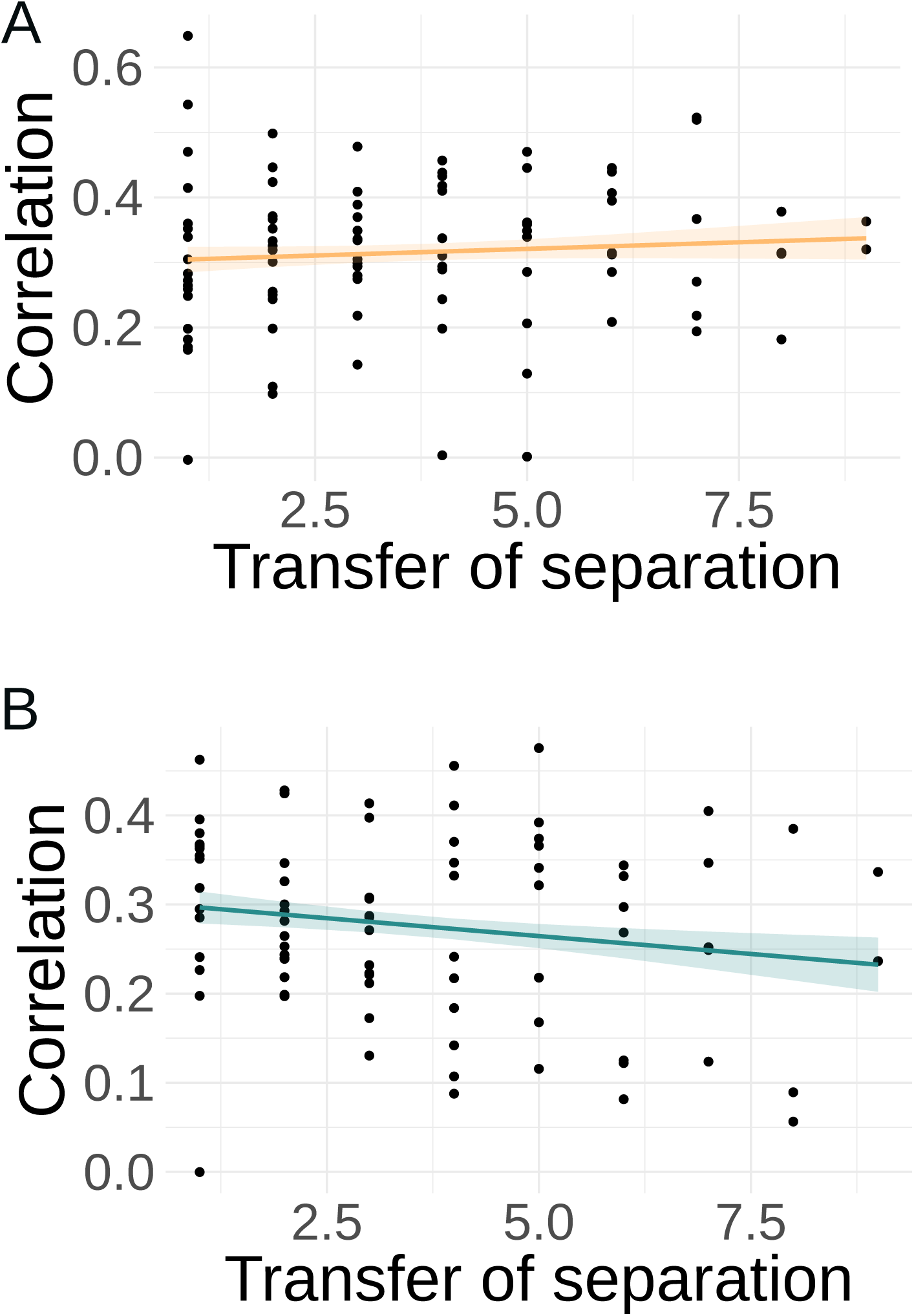
Divergence of the epigenetic changes in typical heterochromatin regions (centromeres and telomeres) in **A**) ethanol and **B**) control.

**Figure S3:**
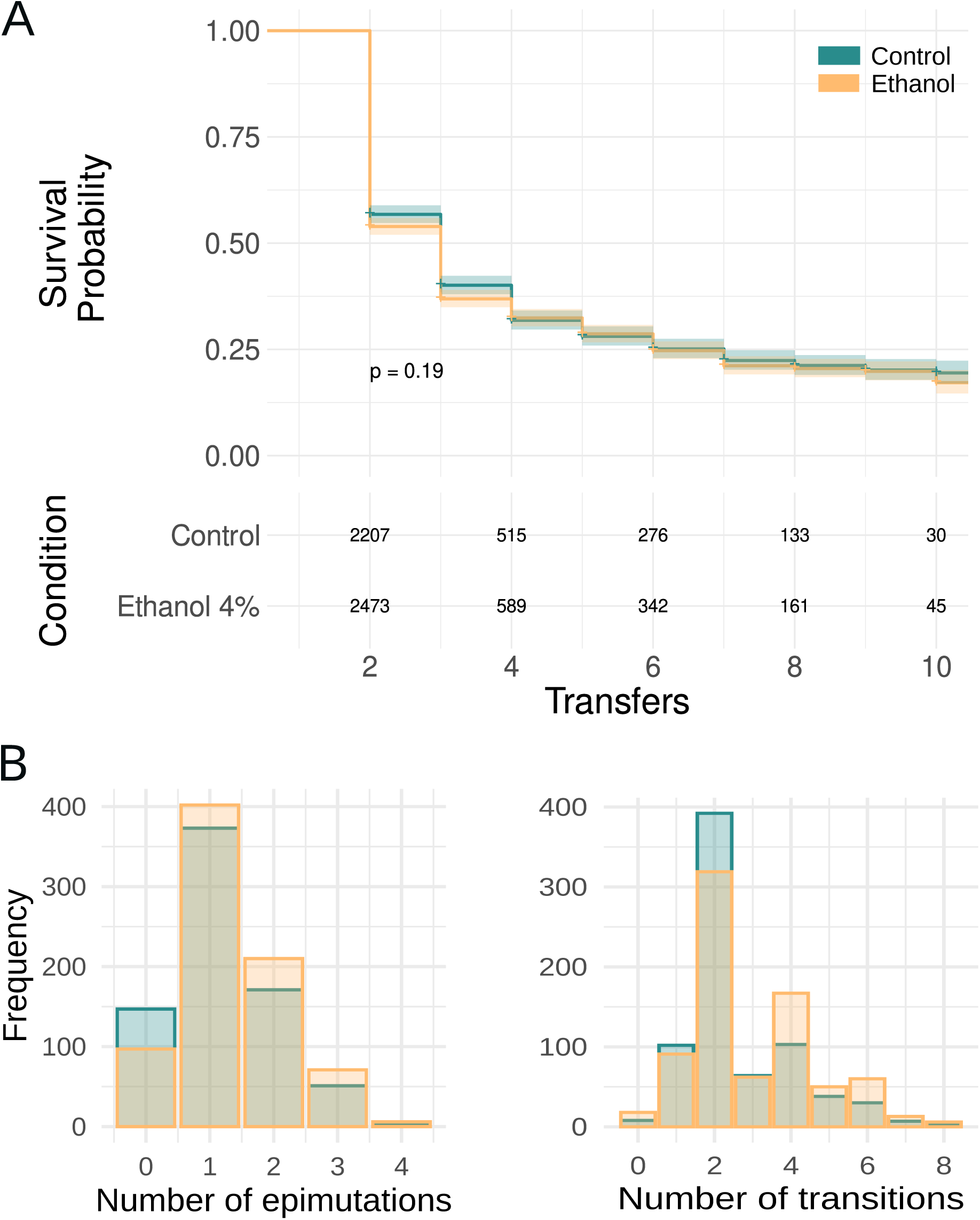
**A**) Survival probability plot and table showing the number of epimutations that are stable across transfers. **B**) Frequency of number of epimutations and transitions

**Figure S4:**
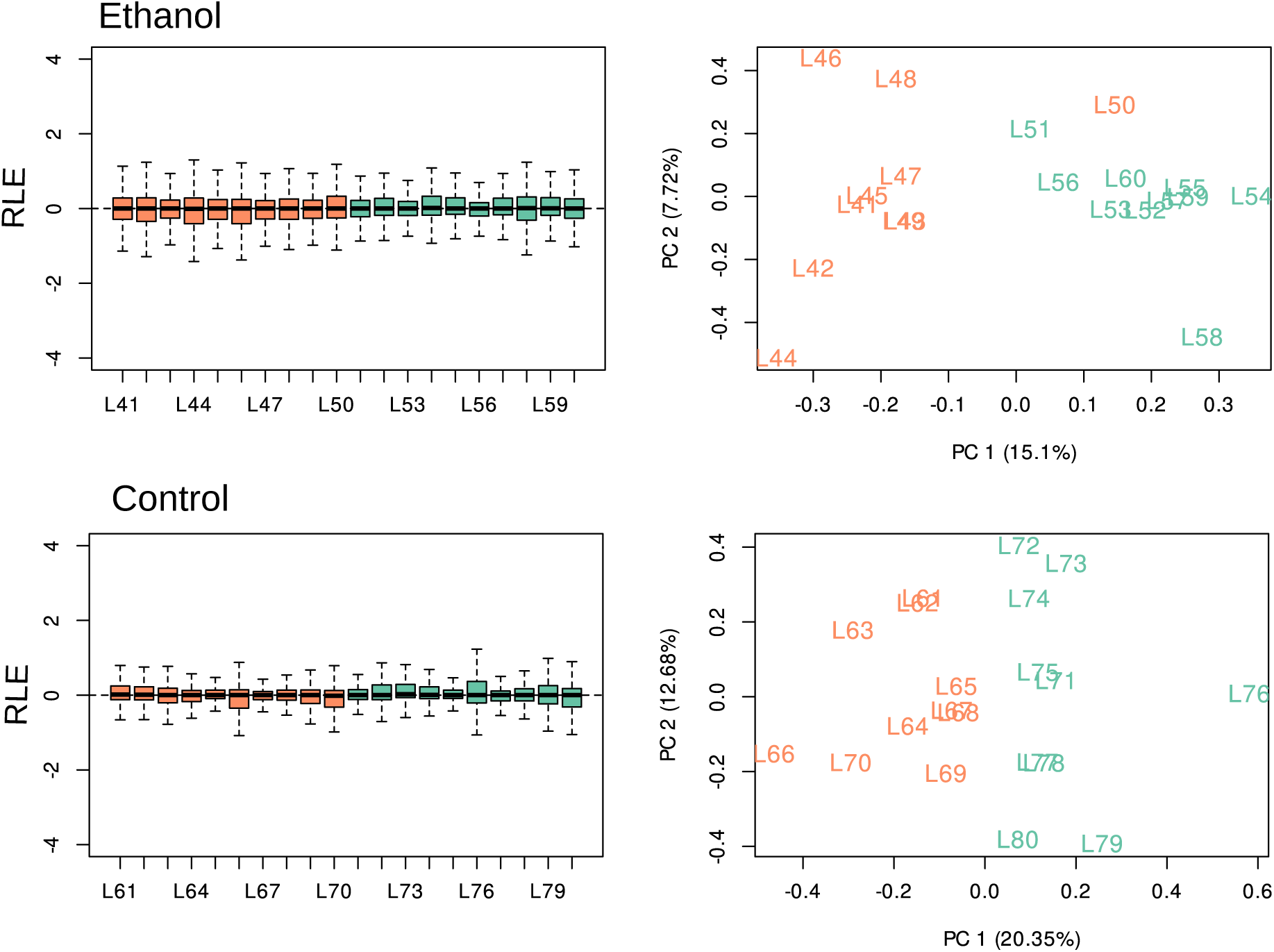
RLE (relative log expression) plots and PCA showing WT and mutant Δago1 RNA-seq data after normalization and removing the unwanted data variation, using *C.elegans* RNA as external spike-in. Colors represent the different strains; in orange the WT and in green Δago1

**Figure S5:**
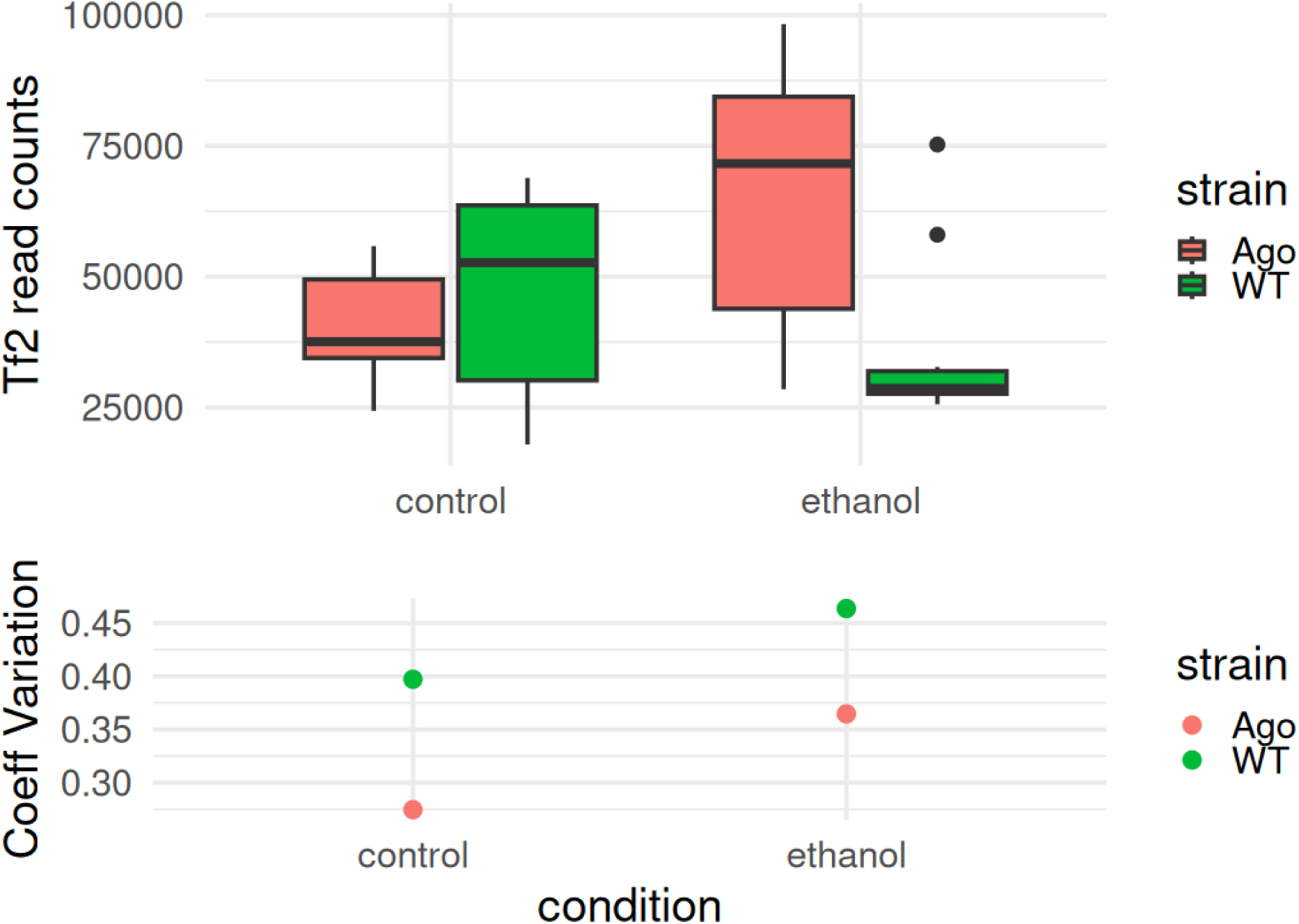
Number of the total counts, and coefficient of variation of reads mapped to the Tf2 family. WT is in pink and Δ*ago1* is in green.

## Supplementary Tables

**Table S1:** Showing the parameters and results for the models that examine number of epimutation, transition, divergence and gene expression noise.

| Model | Response | Fixed-effect | Random factors | Estimate | SE | Stat | p-value | df |
| --- | --- | --- | --- | --- | --- | --- | --- | --- |
|  | Number of epimutations | Treatment + offset(log(number_transfers)) | eMA-line | 0.20 | 0.08698 | z-value = 2.31 | 0.02 | – |
|  | Number of transitions | Treatment + offset(log(number_transfers)) | eMA-line | 0.20 | 0.08698 | z-value = 3.34 | 0.000814 | – |
| Control | Spearman correlation per window | Generation | – | -0.008 | 0.004734 | t-value = -1.801 | 0.07 | 79 |
| Ethanol | Spearman correlation per window | Generation | – | -0.009 | 0.003297 | t-value = -2.779 | 0.00666 | 88 |
| Control heterochromatin | Spearman correlation per window | Generation | – | -0.008023 | 0.005230 | t-value = -1.534 | 0.12 | 79 |
| Ethanol heterochromatin | Spearman correlation per window | Generation | – | 0.004076 | 0.005646 | t-value = 0.722 | 0.47 | 88 |
| Ethanol | log(Coefficient of variation) | Strain | Gene id | 0.26 | 0.00379 | t-value = 68.8 | 2.2e-16 | 6449.00184 |
| Control | log(Coefficient of variation) | Strain | Gene id | 9.173e-02 | 3.976e-03 | t-value = 23.07 | 2.2e-16 | 6.397e+03 |
|  |  | Strain (WT vs mutant) in control | Gene id | 9.173e-02 | 4.325e-03 | t-value = 21.20 | 2e-16 | 1.883e+04 |
| Interaction | log(Coefficient of variation) | Treatment (ethanol vs control) in mutant | Gene id | 2.193e-02 | 4.336e-03 | t-value = 5.05 | 4.27e-07 | 1.883e+04 |
|  |  | Strain × treatment interaction | Gene id | 1.694e-01 | 6.107e-03 | t-value = 27.746 | 2e-16 | 1.883e+04 |

**Table S2:** Table showing the results of the permutation analysis of overlaps between siRNA changes and mRNA changes for TEs.

|  | Observed | Expected | p value | Enrichment |
| --- | --- | --- | --- | --- |
| High mRNA, high siRNA | 0.00617 | 0.0544 | 1 | 0.113 |
| High mRNA, low siRNA | 0.137 | 0.0549 | 0.0001 | 2.49 |
| Low mRNA, high siRNA | 0.0105 | 0.0532 | 1 | 0.198 |
| Low mRNA, low siRNA | 0.407 | 0.0526 | 0.865 | 0.774 |

